# Too many big promises: What is holding back cyanobacterial research and applications?

**DOI:** 10.1101/2023.06.05.543618

**Authors:** Nicolas M. Schmelling, Moritz Bross

**Affiliations:** Krauts & Sprouts, Düsseldorf, Germany; CyanoWorld, Düsseldorf, Germany; Department of Microbiology and Immunology, University of British Columbia, Vancouver, BC, Canada; BASF SE, Ludwigshafen, Germany

**Keywords:** Cyanobacteria, Questionnaire, Challenges, Community

## Abstract

Climate change as a global crisis demands a shift from a fossil fuel-based economy towards sustainable solutions. Cyanobacteria are promising organisms for the truly sustainable, carbon-neutral production of various chemicals. However, so far, proof of concepts for large-scale cyanobacterial productions that produce industrial-relevant amounts of desired products are lacking. To systematically address this topic, a comprehensive overview that identifies current obstacles and solutions is missing.

We conducted a quantitative survey among researchers in the cyanobacterial community. This work investigates individual experiences and challenges in the field of cyanobacteria, as well as information about specific protocols. Additionally, qualitative interviews with academic experts were conducted. Their answers were compared, and highlights were summarised.

In this work, we provide for the first time a comprehensive overview of current trends and challenges as perceived by researchers in the field of cyanobacteria. Based on the results of the survey and interviews, we formulate a set of recommendations on how to improve the working conditions within the cyanobacteria research community.

## 1 Introduction

Cyanobacteria have fascinated the microbiology community for many decades. Unlike any other bacterium, they grow with the help of oxygenic photosynthesis. This allows them to fix carbon from the atmosphere and convert it into complex chemicals that represent valuable products for industrial applications. Hence, many researchers predicted that cyanobacteria will play a significant role in a fossil-free, carbon-neutral industry of the future.

Despite these high expectations, so far, cyanobacteria are only used for the production of specialized compounds, such as fine chemicals or pigments, or as a food supplement in the case of *Spirulina*. In this work, we will shed light on what are the biggest obstacles when it comes to working with cyanobacteria and what researchers, who work with these organisms, consider the main challenges.

## 2 Methods

We combined a quantitative survey with qualitative interviews to get a better understanding of the current challenges within the cyanobacterial research community.

### 2.1 Questionnaire

A comprehensive questionnaire was created and distributed among the members of the cyanobacterial research community to get an overview of the current opinions from within the community.

The questionnaire was distributed via email, Twitter, and through colleagues. The document contained two categories: 1) Scaled questions were used to rate trends in the field. 2) Qualitative questions allowed the participants to provide in-depth answers. We independently categorized the qualitative answers into a set of groups, which were based on the answers given. Afterward, categories were compared and discussed.

### 2.2 Expert interviews

To get in-depth knowledge about the current situation in the cyanobacterial research community, we conducted interviews with renowned experts from the field. Each interview was about one hour in length. All experts we talked to were professors in the field of cyanobacteria. The questions covered three topics: potential products from cyanobacteria, tools, and techniques, and social aspects of the cyanobacterial research community. The most relevant key thoughts from all interviews were extracted, and representing quotes are presented in the results section.

### 2.3 Literature search

We analyzed the methods sections of 349 randomly selected open-access publications working with *Synechocystis* sp. PCC 6803. From this, we extracted the cultivation conditions, like light intensity or growth temperature. These data were also used to validate the representability of the researchers who participated in our questionnaire.

### 2.4 Data Analysis and Availability

The raw and cleaned data from the survey, the data of the literature as well as the Jupyter Notebook analyzing the data are available on figshare (10.6084/m9.figshare.23282879). For analysis of the data, the Python packages, jupyter notebook ([11]), numpy ([5]), matplotlib ([9]), and pandas ([14]) were used.

## 3 Results & Discussion

For this project, we asked 143 researchers within the cyanobacterial research community about their opinion in the field. The participants were mostly male (53 %), while 41 % were female, and 1 % identified as diverse (Fig. 1A). The remaining 5 % preferred not to specify their gender (Fig. 1A). The majority of the participants are located in Europe (114 out of 143), with the rest of them spread among the rest of the world (Fig. 1B). The group of participants consisted of mostly experienced researchers: 28 % worked with cyanobacteria for 2 - 5 years and 18 % for 5 - 10 years (Fig. 1C). The largest portion, 38 %, worked for more than 10 years in the field (Fig. 1C). The vast majority of 90 % are academic researchers (Fig. 1D). Of those, 98 % had at least one university degree, 42 % were postdoctoral researchers or on tenure track positions, and 16 % were full professors (Fig. 1E). The experts we interviewed were all professors working for more than 10 years with cyanobacteria. The results from their interviews are integrated as quotes in the following sections.

**Figure 1:**
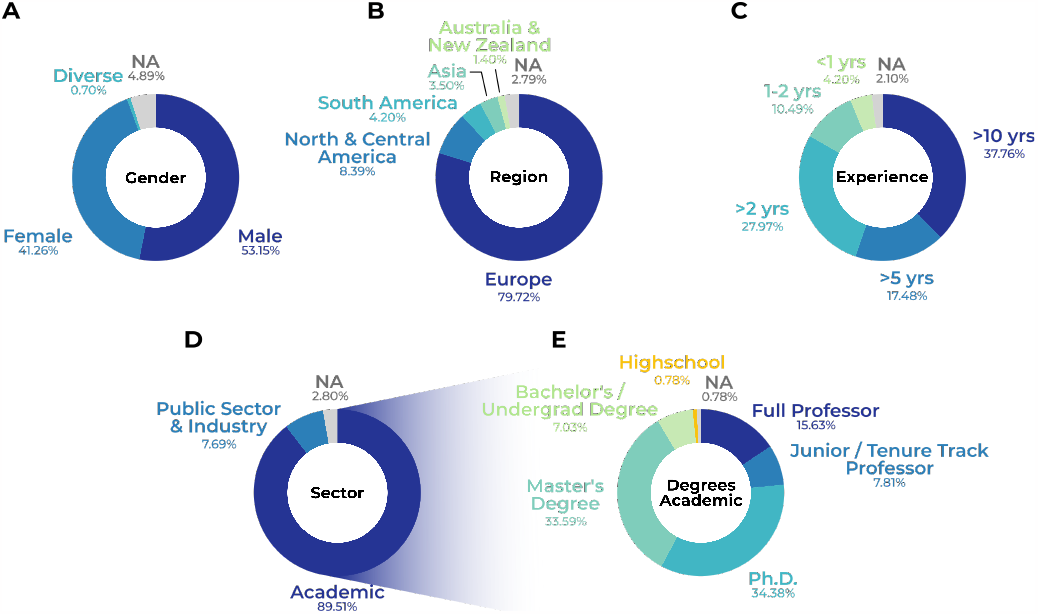
Background information about the participants of this questionnaire. A) gender, B) geographical region, C) experience working with cyanobacteria (years), D) sector of work, E) academic background. N = 143.

### 3.1 Challenges and opportunities of cyanobacterial research

Based on the results of the questionnaire, it is evident that the biggest challenge when working with cyanobacteria appears to be a lack of knowledge about the metabolism of cyanobacteria, including databases and proper annotation (Fig. 2A). As one expert put it: *“Something as simple and basic as genome annotation is still an area where there is opportunity for improvement*.*”* Even once data like genome annotations are created, they are not always properly managed: *“The problem with all these databases is that someone starts them, and then the money runs out, and no one updates them anymore*.*”* A recent initiative CyanoCyc (https://cyanocyc.org/) attempts to end this problem by providing a paid service that curates genomic and metabolic information of cyanobacteria. Another frequently mentioned problem is the lack of standards, such as a widely accepted model strain like *Escherichia coli* K12. Although many researchers work with the strain *Synechocystis* sp. PCC 6803 (34 % of all participants), there are many different substrains used, which sometimes heavily differ in their phenotypes (see Section 3.3.3).

**Figure 2:**
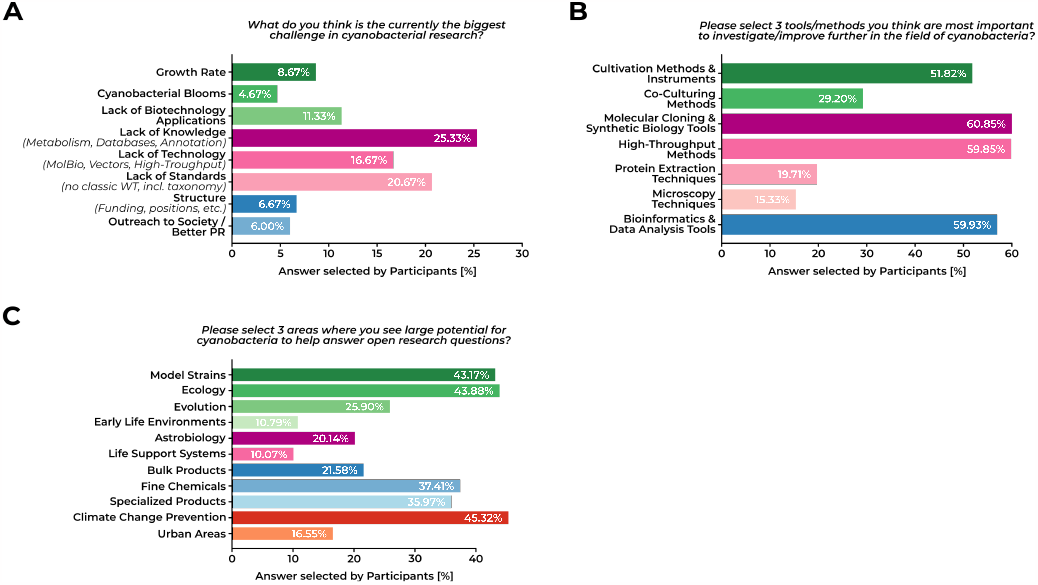
Challenges when working with cyanobacteria. Answers provided by the participants of the questionnaire. A) biggest challenges when working with cyanobacteria (one answer possible, N = 122), B) tools and methods which need improvement (three answers possible, N = 139), C) research areas where work with cyanobacteria could provide answers (up to three answers possible, N = 140).

One survey participant expands on this, saying *“One major problem is the usage of too many different model strains. In the case of Synechocystis sp. PCC 6803 nearly every institute uses their own strain*.*”* We will dive deeper into the reproducibility of cyanobacterial research in Section 3.3.3.

When asked about the specific methods and tools, which are limiting their work, many different aspects were considered hindering (Fig. 2B). More than half of the participants considered molecular cloning, high-throughput methods, bioinformatic tools, and cultivation methods the most pressing aspects (in descending order).

The problem is further described by a comment of another participant: *”Inefficiency in the field of synthetic biology for cyanobacteria: The advent of ‘commercializable’ cyanobacterial biotechnology rather came with more unnecessary competition, ‘every second’ metabolic engineering paper shows another comparison of genetic part performance that is not comparable to other studies (due to lacking standards), claiming to provide another new ‘genetic toolbox for cyanobacterial engineering’. But still, every introduction goes: ‘[*…*] limited selection of genetic tools for cyanobacteria [*…].*’ In fact, the limited tools are just re-iterated from study to study, with inconsistencies between labs*.*”*

The expert interviews revealed further obstacles to cyanobacterial cultivation. For example, light availability within reactor systems is a limiting factor for cyanobacterial growth ([8]). The main issue is the light penetration from outside into a water column (normally sunlight or, alternatively, LED light). While at the outer layers, light intensity is high, inner layers are light-limited ([6, 13]). Different ways to overcome this problem have been proposed. Either by thorough mixing of the culture, cells move between outer and inner layers to get, on average, enough light ([8, 2]). While this creates fluctuating light conditions for the cells, it has been shown that this has no effect on photoinhibition or non-photochemical quenching ([2]). Other alternatives include the reduction of light-harvesting complexes by genetic manipulation ([18]), thin-layer reactor designs, or internal illumination by LEDs ([6]).

When the participants of the questionnaire were asked what they considered challenging about the work with cyanobacteria, they mentioned the slow creation of genetic mutants as well as inconsistencies regarding cyanobacterial taxonomy (including different nomenclatures for genes). Recently, major improvements regarding the cyanobacterial taxonomy and nomenclature have been announced, which will make naming new cyanobacterial strains in the future easier and more reliable ([17]).

Although faced with challenges when working with cyanobacteria, the questionnaire participants saw many fields where cyanobacteria could help answer open research questions (Fig. 2C). Among the top-ranked areas were 1) climate challenge preventions, 2) ecology, and 3) new model strains.

### 3.2 Challenges and opportunities within the cyanobacteria research community

Besides specific problems of working with cyanobacteria, the participants were also asked what they wanted to change about how the cyanobacterial research community works together. In coherence with the data shown in Fig. 2A, the researchers stated that increasing reproducibility and creating standard operating procedures are among their top priorities (Fig. 3A). Furthermore, participants highlighted that the competition within the group, as well as a lack of cooperation, is something that they would like to change in the future.

**Figure 3:**
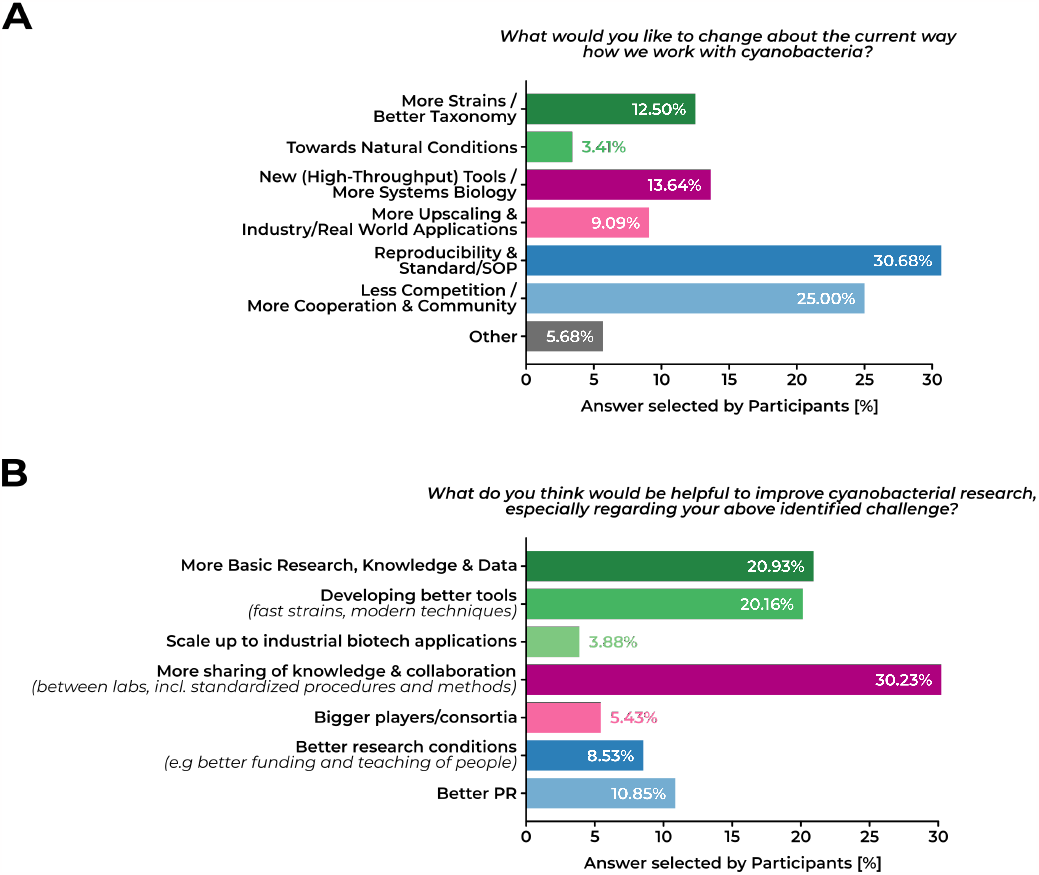
Unique problems within the cyanobacterial research community. A) Suggestions for aspects that should change how the cyanobacterial research community works (N = 88), B) suggested means to achieve the changes mentioned in A) (N = 113).

Consequently, when the participants were asked about improvements to the identified challenges, many stated they would favor more collaboration between labs, including more standardized methods and procedures (Fig. 3B).

Several comments in the open text section mentioned specific measures that could be taken, like *“a standardized, minimized set of information, which has to be reported about culturing analogous to e*.*g. MIQE,”* or *“a website/server, which provides working and verified protocols. Something like www.openwetware*. *org especially for cyanobacteria. Such an encyclopedia would improve reproducibility in the community*.*”* With regard to the latter comment, we would like to highlight the already existing protocol collection of 45 protocols (at the time of writing) specifically tailored to cyanobacteria provided by members of the community on www.protocols.io (https://www.protocols.io/workspaces/cyanoworld). In the future, the list of protocols could be extended or verified to create a set of core protocols, for example, in an event like a hackathon.

Furthermore, an *”index of expertise”* was mentioned. *”A list of institutes and scientists with their expertise would be very useful. [*…*] Next to the technical fields, such a list could also provide information about the research focus*.*”* We would like to highlight the newly created website www.cyano.world, which features such a list (https://www.cyano.world/labs). However, the set-up of this list just started, and available information is still limited at the time of writing, but it is likely to grow in the future.

It also became apparent that larger organized collaborations could improve the effectiveness of several research projects. *”Strengths and resources need to be pooled more. Bigger consortia that actually collaborate with each other in a synergistic fashion, including open data sharing [and] pre-publication. Instead of six labs that each develop their own modular cloning strategy, six labs [should] discuss and develop [a] common strategy (strategies), with comparative studies in each lab*.*”*

Another problem mentioned was the lack of any organized structures which could represent the cyanobacterial community. This could also include sharing certain tasks or the specialization of different working groups, for example, to coordinate tasks like managing databases. The participants suggested a better exchange of knowledge and improved comparability of data. This also includes the publication of negative results or problems within studies.

Lastly, in coherence with the results from Fig. 2A, the researchers also demanded more basic research to get a better understanding of cyanobacteria and the development of better tools like high-throughput methods or model strains.

### 3.3 Lack of reproducibility and standards

From our experience working with cyanobacteria, we anticipated some level of complexity with regard to cultivation, especially considering the large diversity among cyanobacterial strains. The goal of this part of the survey was to identify common themes across cyanobacteria and highlight areas for improving reproducibility. Since “reproducibility” was mentioned many times by the participants, we dedicated an entire section to this topic.

We encountered a plethora of strains used by the participants ranging from marine, over toxic, to soil cyanobacteria, covering a wide range of cyanobacterial species (Fig. 4A). However, six strains stood out: *Synechocystis* sp. PCC 6803 (hereafter Syne6803), *Synechoccocus elongatus* PCC 7942, *Microcystis aeruginosa* NIES-843, *Nostoc* sp. PCC 7120, *Synechococcus* sp. UTEX 2973, and *Synechoccocus* sp. PCC 7002 (Fig. 4A). Those strains are not surprising as they are either the cyanobacterial model strains for their respective class of cyanobacteria or emerging strains. The majority of the participants stated to work with the model strain Syne6803 (Fig. 4A). Due to the amount of data on Syne6803 from this survey, we focused our analysis on those responses to get reliable quantifications. Further, we analyzed a random selection of 349 open-access publications, which involve Syne6803, to get a better understanding of whether our survey data are representative of the cyanobacterial community. We encourage readers to also have a look at the underlying data to find more information on other strains.

**Figure 4:**
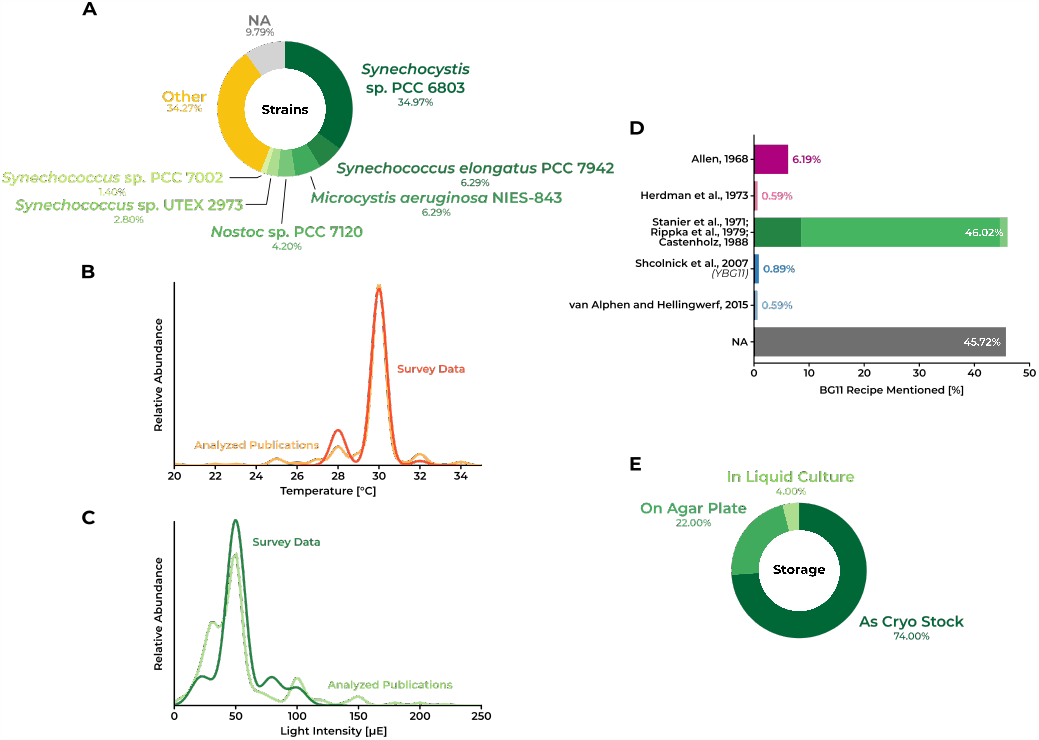
Growth conditions of cyanobacteria. A) Most commonly used cyanobacterial strain, B) temperature used for cultivation, C) light intensity used for cultivation, D) reference used for the BG11 medium most commonly used, E) techniques for the storage of cyanobacteria. The results are based on 143 answers provided by the participants of the questionnaire. Additionally, for B), C), and D), 349 open-access research publications involved work with *Synechocystis* sp. PCC 6803 were screened, and their answers were compared to the ones from the questionnaire.

For Syne6803, several wild-type (WT) strains are known, which can have vastly different phenotypes regarding, among others, their motility or glucose sensitivity ([30, 15]). However, those details about the specific WT strains used in the respective studies are often not mentioned in the final publication. Further, based on personal communication with colleagues and our own experience, we know that, for example, some genes are essential in a certain WT background, whereas in another WT background, they can easily be knocked out. Again, those relevant information are usually also missing in the final publication. Thus, we invite all readers to also include information in the final publication about, for example, a failure of a gene knock-out in one WT strain while successful in another WT strain.

To assess the uniformity within the Syne6803 community, we asked the questionnaire participants to share information about the most common growth parameters, like light intensity, temperature, shaking speed, *CO*_2_ concentration, medium recipes, optical density (OD) measurements, and more. We further analyzed those factors in 349 open-access publications working with Syne6803. First, we find a strong overlap between the cultivation conditions mentioned in those papers and the results of our survey. This indicates that the survey results nicely represent the literature and, thus, the large Syne6803 community, assuming that the randomly selected publications are also a representative selection (Fig. 4B and C).

One would expect some variety within growth parameters, given the just mentioned different WT backgrounds. While we identified two commonly used and very similar growth temperatures of 28°C and 30°C, with a clear preference for 30°C (Fig. 4B), we see large differences among the light intensity. Most strains are cultivated at 50 µE (Fig. 4C). However, the range of light intensities used to routinely grow Syne6803 ranges from as low as 10 µE up to 200 µE (Fig. 4C). We believe that those different light intensities used to grow Syne6803 will have an effect over time, such that presumably identical strains from the same background have differentiated into two different strains by laboratory evolution and adaptation to different light intensities. We hypothesize that if those strains would be analyzed, we would encounter similar phenotypes as for high-light and low-light adapted marine *Prochlorococcus* strains ([3]). However, without thorough analysis, we cannot tell whether those different growth conditions have an effect on the physiology of the cells.

For growing Syne6803, BG11 medium is used in almost all cases. As such, the term “standard BG11” is commonly used. However, when analyzing the publications, several different origins of the BG11 medium are cited. We did not ask the participants to specify the origin of their BG11 medium, so we do not know any specifics here. However, when analyzing the literature data, we find that almost half of the participants either did not specify the origin or use the protocol established by Roger Stanier and colleagues from 1971 ([24]), making the protocol from 1971 the presumably “standard BG11” medium (Fig. 4D). Interestingly, this protocol is identical to Rippka *et al*., 1979 and Castenholz, 1988, and in most instances, Rippka *et al*., 1979 are cited as the origin ([21, 4]) (Fig. 4D). Sometimes even Stanier *et al*., 1979 is cited, which links to Rippka *et al*., 1979 when following the DOI. We do not know if the author’s order has changed over time or if there are different PDF versions available, but as for now, we believe that Stanier *et al*., 1979 is wrongly cited in those cases. When we analyzed the composition of the different BG11 media available, we found that for most parts, they are almost identical with some differences for Allen, 1968 and Herdman *et al*., 1973 and a more recent adaptation by Shcolnick *et al*., 2007, which is known under the name YBG11, and two new compositions by van Alphen and colleagues (2015 and 2018) ([1, 7, 23, 28, 27]; 10.6084/m9.figshare.23282879).

The largest difference among BG11 media is found in the iron composition, which has no effect on the final iron concentration in the medium (10.6084/m9.figshare.23282879). However, we know from personal communications that there are some minor differences between the used chemicals. For example, instead of *K*_2_*HPO*_4_, some labs use *KH*_2_*PO*_4_ or *CoCl*_2_ instead of *Co*(*NO*_3_)_2_. The latter allows for the complete removal of nitrogen from the BG11 medium. Those details are, however, often not mentioned as they might not have a large effect on the overall BG11 medium composition.

Further, the wavelength at which the optical density (OD) is measured differed, with 730 nm and 750 nm being the most widely used options. While measuring OD as a proxy for cell biomass is easy, it is a problematic quantity without additional information like cell count or cell dry weight. It has been shown that OD measurements are very difficult to compare between different labs ([16, 25]). Measuring OD at different wavelengths adds to this problem because it has been shown that even presumably identical Syne6803 strains show differences in the absorption spectra ([15]). Reporting the correct biomass is really important when working, for example, with inducers, because different cell biomass will have an effect on inducer concentration per cell.

Further, we also observed differences in the *CO*_2_ concentration and pH values and buffers used when growing Syne6803. However, in most cases, no additional *CO*_2_ was supplied or mentioned. In addition, we found various pH values of the medium and buffers used. Here, TES (5 different concentrations mentioned) and HEPES (8 different concentrations mentioned) are the most used buffers with a pH of either 7.5 or 8 for Syne6803. One survey participant also mentioned this issue: *”By studying the ‘Method’-section of published research papers from different groups across Germany, I realized many differences in basic methods. For example, so far as I know, the basic recipe of BG11-medium contains NaHCO*_3_ *as a pH buffer. However, many groups use routinely TES or HEPES. Previously, I found a paper in which the authors completely waived the buffer and titrated the BG11-medium directly with NaOH and HCl to a pH of 8*.*0. The usage of different model strains coupled with deviating methods decreases reproducibility*.*”*

Lastly, most of the participants mentioned that they store their strains as cyro-stocks (Fig. 4E). However, almost one quarter did use other methods. Most of those stored their strains on agar plates, which is, in our opinion, not a reliable technique for the long-term storage of Syne6803 cells. While you might select certain cells by unthawing your cryo-stock, those cells should have identical genotypes, and selection is based on cell viability rather than genotype. Whereas on agar plates, mutations occur faster than in cryo-stocks, even though slower than in liquid culture, and the likelihood for selection based on different genotypes is higher from restreaking. We acknowledge that cryo-preservation is impossible for many cyanobacterial species. However, for the analyzed Syne6803 it is. We would like to highlight that bacteria mutate fast, and storing them in liquid culture is probably not ideal. It has been shown that “strains can rapidly deviate from their original genotype, which can sometimes lead to noticeable phenotypic differences” and that precautions like cryo-preservation of background strains can mitigate those problems ([19]).

### 3.4 Challenges and potential of cyanobacteria for industrial applications

Since cyanobacteria are still behind their expectations regarding their potential for sustainable production of industrial goods, we asked the survey and interviewed participants more specifically about this topic.

When asked about challenges for the industrial usage of cyanobacteria, participants selected large-scale cultivation as the most limiting factor (Fig. 5A). One of the reasons why it is difficult to upscale cyanobacteria cultures has been described by one expert as the following: *”Probably the biggest issue [of upscaling cyanobacterial production] is contamination. As soon as you scale up, you cannot prevent contamination*.*”* At the same time, one expert stated that the cultivation of cyanobacteria always results in the dilemma that more axenic growth conditions also increase the cultivation costs. So far, we are still lacking a tolerant production strain that withstands contaminations, as well as high temperatures, light intensity, and salt concentrations. Even though there have been advantages in all individual fields ([29, 12]), a robust strain comparable to the production strain *E. coli* is still lacking. The recent development of new high-density cultivation systems, for example, by the German Start-Up CellDEG (https://celldeg.com/), can help to unlock the potential of already existing strains like Syne6803.

**Figure 5:**
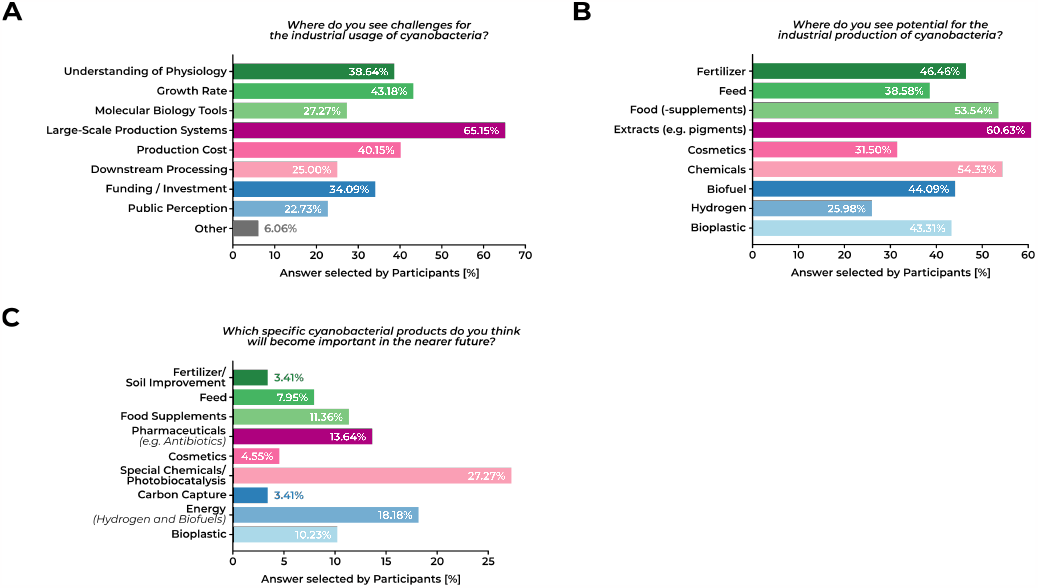
Industrial applications of cyanobacteria. A) Challenges for industrial usage of cyanobacteria (N = 132), B) potential use (N = 127), and C) specific products derived from cyanobacteria (N = 79).

The lack of proper upscaling results in relatively high prices compared to heterotrophic bacteria: *”The competition with heterotrophs will always be unfair unless we are able to charge a premium for the fact that it is a CO*_2_*-based product”* as one expert stated.

*”Understanding where we can compete with current production costs can point us to what we can produce with this system. That’s what happened in the field of biocatalysis, where there was a shift from bulk chemicals to fine chemicals, and that’s the same what happened with bioplastic production, where there was a shift from producing plastic bags (super cheap) to implants (high-cost production)*.*”* Recent life cycle assessments and techno-economic analyses demonstrated that the current production costs for cyanobacteria, exemplified for the bioplastic polyhydroxy butyrate (PHB), are not economically viable ([20, 22]). However, they also highlight strategies to reach profitability. Major drivers of production costs are product yield, equipment costs, and labor and maintenance costs ([22]). First, production yield could be improved through genetic modifications ([22]). As most costs are fixed, the costs per product unit will be significantly reduced by increasing product yield. Next, the selection and costs of the reactor system influence the production costs, especially the volume-to-surface ratio, which influences the productivity of phototrophs, is a major factor ([22]). Last, biorefinery approaches that produce more than one product from the cell biomass, like PHB and pigments, are a solution to make cyanobacterial production economically viable ([20]). However, those concepts are still understudied. Depending on those factors, the break-even minimum selling price, in the case of PHB, can be decreased from 449€/kg down to 7.7€/kg, while the current PHB market price is at 4€/kg ([20, 22]). Similar results are probably obtained for other bulk chemicals like polypropylene/polyethylene, polylactic acid, or ethanol.

While we agree with the experts’ opinions, we believe that higher prices might not be necessary if the true cost of production (including the environmental damage caused by *CO*_2_ emissions) is included in the price of competing products. While it would be a fast intervention to charge a premium to the consumers, this would not be a sustainable, long-term solution. In the long run, it will be necessary to implement substantial regulatory changes, such as higher *CO*_2_ taxes, which would consequently favor cyanobacterial products.

Furthermore, another expert stated that *”GMO acceptability is still an obstacle”* when it comes to commercializing cyanobacterial products. Hence it is crucial to have a societal discussion about this topic since *”we cannot really do anything without genetic engineering*.*”*

In addition, we believe that a lack of GRAS (Generally Recognized As Safe) status for cyanobacteria is an obstacle. So far, only *Spirulina* is considered safe for human consumption. However, almost all molecular tools are developed for other cyanobacterial strains since working with *Spirulina* has been challenging. Recently, researchers from Lumen Bioscience (https://www.lumen.bio/) managed to genetically modify *Spirulina* and published the protocol ([10, 26]). We believe that this will facilitate the adoption of cyanobacteria as production hosts. Furthermore, we believe that the increase of cyanobacterial strains with GRAS status will further boost the biotechnological adaptation of these microorganisms.

When asked about potential compounds, which may be produced in cyanobacteria, the answers were widespread (Fig. 5B). The most frequently given answers were pigments, chemicals, and food supplements, which is in line with currently available products from *Spirulina* production. One expert mentioned that the potential of cyanobacteria to fix nitrogen might be exploited for food production. At the same time, another expert highlighted that it is difficult to produce well-tasting cyanobacteria due to their own natural taste.

Further products mentioned were special chemicals/products of photocatalysis (Fig. 5C). Additionally, the usage of cyanobacteria for energy carriers, such as hydrogen or biofuels, was mentioned. Despite the wide field of potential applications, several experts stated that the cyanobacterial community still lacks a sense of entrepreneurship. As one expert said, *”maybe what we need is also an example of ‘yes, we can do it’*.*”* He also mentioned that, in order to make cyanobacteria more successful, *”we need the influencers, we need pioneers, we need leading countries*.*”*

### 3.5 Conclusion

Our analysis revealed that there are large variations in how different research groups work with cyanobacteria. Hence, establishing standards for the entire community would be a crucial step to improve the quality of data and hypotheses generated.

Based on the reproducibility analysis of Syne6803, it would be, as one participant mentioned, *”extremely useful if the [*…*] community could manage to agree on one”* strain. Further, we like to encourage everyone to make cryo stocks from their Syne6803 strains and use a fresh batch for every important set of experiments. As for growth conditions, we propose to use 50 µE and 30°C and the BG11 medium from Stanier *et al*., 1971 without any additional buffer. Feel free to refer to this composition as the ”standard BG11.” We encourage you to measure the light spectra of your incubator or get the light specifications from the manufacturer as a CSV file and share it in the supplement of your future publications. Keep using OD measurements (at least at 730 nm) as a proxy for cell biomass, as it is easy and reliable within a lab. However, we propose making calibration curves with cell counts at least once a year to have this additional information ready to share with your publication. Applying those measures will increase the reproducibility of Syne6803 research substantially. Not all of those suggestions apply to other cyanobacterial strains, but we encourage you to think about increasing the reproducibility of your strain as best as possible.

In accordance with the opinion of our experts and the wide cyanobacteria research community, we would encourage the founding of more start-ups and industry collaboration. This would leverage the upscaling of cyanobacterial cultivation, which is another major limiting factor regarding their commercialization. Finally, more political support is necessary, for example, regarding regulating GMOs, to unleash the full potential of cyanobacterial biotechnology.

While there are still many obstacles when it comes to the application of cyanobacteria, we see the potential for long-term projects, assuming the proposed measures are implemented.

## 4 Conclusion

This survey provides valuable insights into the challenges and opportunities within the cyanobacteria research community. It is evident that establishing supporting infrastructure, such as a maintained database for omics data and laboratory protocols, is crucial. Further, the community highlighted their wish for more collaboration and larger research consortia over fragmented projects. However, connections between laboratories need to be addressed to improve efficiency.

Standardizing research practices and establishing guidelines for working with cyanobacteria would greatly enhance data quality and reproducibility. Based on the reproducibility analysis of Syne6803, it would be, as one participant mentioned, *“extremely useful if the [*…*] community could manage to agree on one”* strain for Syne6803. Further, we like to encourage everyone to make cryo stocks from their Syne6803 strains and use a fresh batch for every important set of experiments. As for growth conditions, we propose to use 50 µE and 30°C and the BG11 medium from Stanier et al., 1971 without any additional buffer as “standard” growth conditions. We encourage you to measure the light spectra of your incubator or get the light specifications from the manufacturer as a CSV file and share it in the supplement of your future publications. Keep using OD measurements (at least at 730 nm) as a proxy for cell biomass, as it is easy and reliable within a lab. However, we propose making calibration curves with cell counts at least once a year to have this additional information ready to share with your publication. Applying those measures will increase the reproducibility of Syne6803 research substantially. Not all of those suggestions apply to other cyanobacterial strains, but we encourage you to think about increasing the reproducibility of your strain as best as possible.

The cyanobacteria research community recognizes the challenges in utilizing cyanobacteria as a biotechnology production platform. However, they also acknowledge the immense potential once these challenges are overcome. Encouraging the establishment of start-ups and fostering industry collaboration can facilitate the scaling of cyanobacterial cultivation. Additionally, political support is crucial, particularly in regulating GMOs, to fully unlock the potential of cyanobacterial biotechnology.

Although obstacles remain, long-term projects hold promise if the proposed measures are implemented effectively.

## Author Contributions

M. B. and N. M. S. both conducted the interviews, designed the questionnaire, interpreted the data, designed the figures, and wrote the original manuscript.

## Acknowledgments

The authors would like to thank all experts who took the time to participate in the interview process. We also want to thank all survey participants for their contribution.

## References

[1] M. M. Allen. Simple conditions for growth of unicellular blue-green algae on plates. Journal of Phycology, 4(1):1–4, 1968.

[2] B. Andersson, C. Shen, M. Cantrell, D. S. Dandy, and G. Peers. The fluctuating cell-specific light environment and its effects on cyanobacterial physiology. Plant Physiology, 181(2):547–564, 2019.

[3] S. J. Biller, P. M. Berube, D. Lindell, and S. W. Chisholm. Prochlorococcus: the structure and function of collective diversity. Nature Reviews Microbiology, 13(1):13–27, 2015.

[4] R. W. Castenholz. Culturing methods for cyanobacteria. In Cyanobacteria, volume 167 of Methods in Enzymology, pages 68–93. Academic Press, 1988.

[5] C. R. Harris, K. J. Millman, S. J. van der Walt, R. Gommers, P. Virtanen, D. Cournapeau, E. Wieser, J. Taylor, S. Berg, N. J. Smith, R. Kern, M. Picus, S. Hoyer, M. H. van Kerkwijk, M. Brett, A. Haldane, J. F. del Río, M. Wiebe, P. Peterson, P. Gérard-Marchant, K. Sheppard, T. Reddy, W. Weckesser, H. Abbasi, C. Gohlke, and T. E. Oliphant. Array programming with numpy. Nature, 585(7825):357–362, 2020.

[6] M. Heining, A. Sutor, S. C. Stute, C. P. Lindenberger, and R. Buchholz. Internal illumination of photobioreactors via wireless light emitters: a proof of concept. Journal of Applied Phycology, 27(1):59–66, 2015.

[7] M. Herdman, S. F. Delaney, and N. G. Carr. A new medium for the isolation and growth of auxotrophic mutants of the blue-green alga Anacystis nidulans. Microbiology, 79(2):233–237, 1973.

[8] Q. Huang, F. Jiang, L. Wang, and C. Yang. Design of photobioreactors for mass cultivation of photosynthetic organisms. Engineering, 3(3):318–329, 2017.

[9] J. D. Hunter. Matplotlib: A 2d graphics environment. Computing in Science Engineering, 9(3):90–95, 2007.

[10] B. W. Jester, H. Zhao, M. Gewe, T. Adame, L. Perruzza, D. T. Bolick, J. Agosti, N. Khuong, R. Kuestner, C. Gamble, K. Cruickshank, J. Ferrara, R. Lim, T. Paddock, C. Brady, S. Ertel, M. Zhang, A. Pollock, J. Lee, J. Xiong, M. Tasch, T. Saveria, D. Doughty, J. Marshall Carrieri, L. Goetsch, J. Dang, N. Sanjaya, D. Fletcher, A. Martinez, B. Kadis, K. Sigmar Afreen, T. Nguyen, A. Randolph, A. Taber, A. Krzeszowski, B. Robinett, D. B. Volkin Grassi, R. Guerrant, R. Takeuchi, B. Finrow, C. Behnke, and J. Roberts. Development of Spirulina for the manufacture and oral delivery of protein therapeutics. Nature Biotechnology, 40(6):956–964, 2022.

[11] T. Kluyver, B. Ragan-Kelley, F. Pérez, B. Granger, M. Bussonnier, J. Frederic, K. Kelley, J. Hamrick, J. Grout, S. Corlay, P. Ivanov, D. Avila, S. Abdalla, C. Willing, and Jupyter development team. Jupyter notebooks - a publishing format for reproducible computational workflows. In F. Loizides and B. Scmidt, editors, Positioning and Power in Academic Publishing: Players, Agents and Agendas, pages 87–90. IOS Press, 2016.

[12] M. Koch, A. J. C. Noonan, Y. Qiu, K. Dofher, B. Kieft, S. Mottahedeh, M. Shastri, and S. J. Hallam. The survivor strain: isolation and characterization of Phormidium yuhuli ab48, a filamentous phototactic cyanobacterium with biotechnological potential. Frontiers in Bio-engineering and Biotechnology, 10, 2022.

[13] B. F. Lucker, C. C. Hall, R. Zegarac, and D. M. Kramer. The environmental photobioreactor (epbr): An algal culturing platform for simulating dynamic natural environments. Algal Research, 6:242–249, 2014.

[14] W. McKinney. Data Structures for Statistical Computing in Python. In S. van der Walt and J. Millman, editors, Proceedings of the 9th Python in Science Conference, pages 56–61, 2010.

[15] J. N. Morris, J. J. Eaton-Rye, and T. C. Summerfield. Phenotypic variation in wild-type substrains of the model cyanobacterium Synechocystis sp. PCC 6803. New Zealand Journal of Botany, 55(1):25–35, 2017.

[16] J. A. Myers, B. S. Curtis, and W. R. Curtis. Improving accuracy of cell and chromophore concentration measurements using optical density. BMC Biophysics, 6(1):4, 2013.

[17] A. Oren, D. R. Arahal, R. Rosselló-Móra, I. C. Sutcliffe, and E. R. B. Moore. Emendation of general consideration 5 and rules 18a, 24a and 30 of the international code of nomenclature of prokaryotes to resolve the status of the cyanobacteria in the prokaryotic nomenclature. International Journal of Systematic and Evolutionary Microbiology, 71(8), 2021.

[18] J. E. Polle, S. Kanakagiri, E. Jin, T. Masuda, and A. Melis. Truncated chlorophyll antenna size of the photosystems — a practical method to improve microalgal productivity and hydrogen production in mass culture. International Journal of Hydrogen Energy, 27(11):1257–1264, 2002.

[19] C. E. Price, F. Branco dos Santos, A. Hesseling, J. J. Uusitalo, H. Bachmann, V. Benavente, A. Goel, J. Berkhout, F. J. Bruggeman, S.-J. Marrink, M. Montalban-Lopez, A. de Jong, J. Kok, D. Molenaar, B. Poolman, B. Teusink, and O. P. Kuipers. Adaption to glucose limitation is modulated by the pleotropic regulator ccpa, independent of selection pressure strength. BMC Evolutionary Biology, 19(1):15, 2019.

[20] S. Price, U. Kuzhiumparambil, M. Pernice, and P. Ralph. Techno-economic analysis of cyanobacterial phb bioplastic production. Journal of Environmental Chemical Engineering, 10(3):107502, 2022.

[21] R. Rippka, J. Deruelles, J. B. Waterbury, M. Herdman, and R. Y. Stanier. Generic assignments, strain histories and properties of pure cultures of cyanobacteria. Microbiology, 111(1):1–61, 1979.

[22] E. Rueda, V. Senatore, T. Zarra, V. Naddeo, J. García, and M. Garfí. Life cycle assessment and economic analysis of bioplastics production from cyanobacteria. Sustainable Materials and Technologies, 35:e00579, 2023.

[23] S. Shcolnick, Y. Shaked, and N. Keren. A role for mrga, a dps family protein, in the internal transport of fe in the cyanobacterium Synechocystis sp. PCC 6803. Biochimica et Biophysica Acta (BBA) - Bioenergetics, 1767(6):814–819, 2007.

[24] R. Y. Stanier, K. R., M. M., and C.-B. G. Purification and properties of unicellular blue-green algae (order chroococcales). Bacteriological Reviews, 35(2):171–205, 1971.

[25] K. Stevenson, A. F. McVey, I. B. N. Clark, P. S. Swain, and T. Pilizota. General calibration of microbial growth in microplate readers. Scientific Reports, 6(1):38828, 2016.

[26] H. Tabakh, B. W. Jester, H. Zhao, R. Kuestner, N. Khuong, C. Shanitta, R. Takeuchi, and J. Roberts. Protocol for the transformation and engineering of edible algae Arthrospira platensis to generate heterologous protein-expressing strains. STAR Protocols, 4(1):102087, 2023.

[27] P. van Alphen, H. Abedini Najafabadi, F. Branco dos Santos, and K. J. Hellingwerf. Increasing the photoautotrophic growth rate of Synechocystis sp. PCC 6803 by identifying the limitations of its cultivation. Biotechnology Journal, 13(8):1700764, 2018.

[28] P. van Alphen and K. J. Hellingwerf. Sustained circadian rhythms in continuous light in Synechocystis sp. PCC 6803 growing in a well-controlled photobioreactor. PLOS ONE, 10(6):1–12, 2015.

[29] A. Wlodarczyk, T. T. Selão, B. Norling, and P. J. Nixon. Newly discovered Synechococcus sp. PCC 11901 is a robust cyanobacterial strain for high biomass production. Communications Biology, 3(1), 2020.

[30] T. Zavřrel, P. čcenášsová, and J. Červený. Phenotypic characterization of Synechocystis sp. PCC 6803 substrains reveals differences in sensitivity to abiotic stress. PLOS ONE, 12(12):1–21, 12 2017.

